# Electron microscopic evidence that Aip1 disintegrates cofilin-saturated F-actin domains in the presence of coronin

**DOI:** 10.1101/199463

**Authors:** Vivian W. Tang, Ambika V. Nadkarni, William M. Brieher

## Abstract

Cofilin is an essential actin filament severing protein necessary for fast actin turnover dynamics. Segments of actin bound to cofilin adapt an alternative twist. This configuration is stable, but boundaries between cofilin occupied and unoccupied polymer are weak and fragment. Coronin and Aip1 are two factors that promote cofilin mediated actin filament disassembly, but whether they simply accelerate the basic cofilin severing mechanism or alter the mode of filament disassembly is still being investigated. Using electron microscopy and spectroscopy, we show that coronin accelerates phosphate release from F-actin to stimulate highly cooperative cofilin binding on to the polymer creating long stretches with a hypertwisted morphology. We find that Aip1 attacks these hypertwisted regions along their length, not just the boundaries, causing sections to disintegrate into monomers. Therefore, coronin promotes cofilin binding to F-actin to generate longer segments of polymer that are themselves the substrates for Aip1 mediated disintegration, as opposed to simply creating more heterotypic junctions that would sever. The morphological characteristics of the disassembling filaments along with spectroscopic data showing the rapid liberation of actin monomers suggest that the combination of cofilin, coronin, and Aip1 might be triggering a more catastrophic mode of filament disassembly than severing.

## Introduction

The rapid disassembly of actin filaments is necessary for the dramatic shape changes that accompany fundamental cellular processes such as cell division, endocytosis, cell motility and wound-healing. Cofilin is an actin filament severing protein that is necessary for fast actin disassembly in cells (1). It binds to F-actin cooperatively (2-8) and alters filament twist (6,9) and filament mechanics (10,11). This alternative twist is stable, but junctions between occupied and unoccupied polymer are weak causing filaments to sever at or near the junction to produce two intact filaments (8,12-15). Therefore, actin filaments are most susceptible to cofilin-mediated severing at intermediate occupancies of cofilin that maximize the number of heterotypic junctions susceptible to fragmentation (16-18).

Given this reaction mechanism, cooperative binding of cofilin seemingly would be counterproductive to cofilin’s primary function of disassembling actin filaments and promoting fast actin turnover. If severing events required several contiguous cofilins, then cooperative binding might promote filament fragmentation. Live imaging suggests that severing events might need 20 – 100 nanometers of contiguous cofilins (8,14). However, even low concentrations of cofilin are sufficient to sever actin (16,19). Cofilin’s cooperative binding is too weak to produce long stretches of contiguous occupied sites at these low cofilin concentrations which implies that only one or two bound cofilin molecules are necessary for filament fragmentation (3,16,19). What then is the purpose of cooperative binding? Cofilin-saturated domains could be stable until some factor displaced cofilin to create a heterotypic junction that would sever (17). In a major turn of events, Wioland et al showed that cofilin saturated barbed ends are hard to elongate or cap and tend to depolymerize. Thus, cooperative binding helps cofilin reach capped barbed ends, uncap them, and depolymerize the filament from the barbed end (8). A third possibility is that auxiliary factors that enhance cofilin’s activity target cofilin-saturated domains, and not just the junctions (18,20-23).

Cofilin mediated actin disassembly is coupled to ATP hydrolysis on actin filaments. While ATP hydrolysis occurs soon after an actin monomer incorporates into a filament (24), subsequent Pi release is slow and occurs in minutes (25-27). Cofilin binds selectively to ADP-actin and must wait for Pi release before it can act (7,24,28). Pi release is therefore thought to act as a timer for selectively marking old filaments for cofilin mediated disassembly. However, many actin networks in cells, like lamellipodia or Listeria actin comet tails, are highly dynamic and turn over on time scales under a minute (29-32). Factors must exist that accelerate Pi release from the actin filament to accelerate depolymerization. Cofilin itself accelerates Pi release (25) but other factors might contribute to the reaction.

Coronin and Aip1 are two factors that can act in concert with cofilin to rapidly disassemble *Listeria* actin comet tails(33) and actin filaments *in vitro* (30,34). Coronin is an F-actin binding protein(35) that promotes cofilin binding to *Listeria* actin comet tails (33) and to F-actin (33,34) and enhances filament severing depending on nucleotide state (36,37). Aip1 accelerates cofilin mediated filament severing (18,23,38) and subunit dissociation from filament ends (18) to produce actin monomer more rapidly than cofilin alone (13,39). Interestingly, Aip1 works most efficiently when cofilin binding densities on F-actin are saturating (18). This suggests that Aip1 can form a ternary complex with cofilin and F-actin (21-23,38) to destabilize sections of cofilin decorated polymer all along their length and not just at heterotypic junctions. How coronin promotes cofilin binding to F-actin is not known (40,41).

Previous results using high speed fluorescence imaging suggested that the combination of cofilin, coronin, and Aip1 triggered a cooperative form of filament disassembly that was referred to as bursting (30). The bursting hypothesis proposed that sections of F-actin would undergo a catastrophic conversion from polymer to monomers and very short oligomers. Catastrophic filament disassembly predicts a more severe disruption of filament architecture than severing, which is orderly and produces two recognizable daughter filaments. Here, we used electron microscopy and spectroscopy to further examine the changes in F-actin that accompany its disassembly in the presence of cofilin, coronin, and Aip1 to see 1) if Aip1 targets sections of F-actin coated with cofilin for disassembly or only heterotypic junctions, 2) how coronin promotes cofilin binding to F-actin, and 3) if this triple mix cocktail of depolymerizers induces a mode of disassembly that is morphologically distinct from severing.

## Experimental Procedures

### FRET assay

Aliquots of labeled actin were diluted to 20 μM in G buffer (pH 7.4) and spun the next day at 227,900 X g for 20 minutes. 35-40% Tetramethylrhodamine labelled actin and 12-15% Oregon green 488 labelled actin were premixed. For depolymerization reactions in the presence of the bursting factors actin was prepolymerized at 10 μM by the addition of 1x F-buffer(10 mM HEPES 7.8, 50 mM KCl, 0.5 mM EGTA, 1 mM MgCl2, 1 mM ATP). The final concentration of actin in the reaction was 1 μM. Spectroscopic monitoring of fluorescence of OG488 (λEx = 490 nm, λEm = 530 nm) over time was used to report on disassembly on a Spectramax M2 fluorimeter (Molecular Devices). Final concentrations of disassembly proteins were 1.25 μM cofilin, 0.75 μM coronin and 0.1 μM Aip1, unless indicated otherwise in graphs. Data was normalized using values of 1 as control actin polymer (in 1xF buffer) and 0 as control actin monomer (in G-buffer).

### Monomer generation assay

Actin for this assay was prepared identically as the FRET assay except for the exclusion of TMR-actin from the reaction. TMR-actin was replaced by unlabelled G-actin. 1 μM Vitamin D-binding protein (Athens Research and Technology) was added to monomeric or polymeric actin to sequester actin monomer and the fluorescence was monitored (λEx = 490 nm, λEm = 530 nm) over time on a Spectramax M2 fluorimeter (Molecular Devices). Concentrations of proteins were as follows: 1 μM actin 1.25 μM cofilin, 0.75 μM coronin 0.1 μM Aip1 and 0.1 μM CapZ when applicable.

### Seeding assay

Actin filaments treated with various combinations of depolymerizers as described in the FRET assay. The reaction was allowed to proceed to completion and 1/10 of the reaction (125 nM) was used to seed new pyrene G-actin assembly (25% labelled, 1 μM, preincubated in G-buffer). Fluorescence was monitored (λEx = 365 nm, λEm = 410 nm) over time.

### Phosphate Release

Phosphate release from F-actin was measured using purine nucleoside phosphorylase described by Webb (42). Briefly, 20 uM G•actin was polymerized in the presence of varying concetrations of cofilin and coronin described in the text in reaction buffer (20mM Tris pH7.5, 50 mM KCl, 2mM MgCl2, 1mM EGTA, 1mM ATP, 0.2 mM methylthioguanosine, and 1 unit of purine nucleoside phosphorylase). Phosphate release was monitored by measuring the increase in optical density at 360 nm using a Spectramax M2 plate reader. The OD 360nm signal was corrected for scattered light from actin polymerization by subtracting the signal obtained at 60 nm in the absence of purine nucleoside phosphorylase.

### Negative Stain Electron Microscopy

To measure actin filament crossover distances, 10 uM G•actin was copolymerized with 12 uM cofilin +/- 2 μM coronin for 90 seconds. Samples were fixed for 30 seconds with 0.1% glutaraldheyde, transferred to glow-discharged, carbon-coated EM grids for 10 seconds and then washed two times in polymerization buffer before staining with 2% uranyl acetate. To monitor filament disassembly in presence of Aip1, actin was copolymerized with cofilin and coronin using concentrations indicated in the results. Aip1 was then injected into the sample to a final concentration of 0.2 μM before fixing with glutaraldehyde at various times thereafter as described above. Electron micrographs were acquired using Tecnai G^2^ Spirit BioTWIN electron microscope at 120kV.

## Results

Cofilin-mediated severing reactions produce two intact actin filaments with no immediate loss in total actin polymer mass. In contrast, the triple mix of cofilin, coronin, and Aip1 is hypothesized to produce actin monomers and very small oligomers with an immediate loss of actin polymer mass. These two alternative models of filament disassembly can be distinguished by imaging actin filaments in the electron microscope before and shortly after adding actin disassembly factors to pre-formed actin filaments. Our basic protocol was to polymerize a solution of actin for a brief period time, add the disassembly factors, fix the reaction with glutaraldehyde, blot the sample onto glow discharged EM grids, stain with uranyl acetate, and image the sample in the electron microscope.

Figure 1 shows an overview of actin filaments before and after treating them with various combinations of cofilin, coronin, and Aip1. Figure 1A is a low magnification view of a 10μM solution of actin that was induced to polymerize for one minute prior to fixation. Note that these are young filaments, and the reaction is still in the rapid assembly phase. Figure 1B shows a representative field when a 10μM solution of actin is polymerized for one minute and then subsequently treated with 5μM cofilin, 2 μM coronin, and 0.2μM Aip1 for an additional sixty seconds prior to fixing and visualizing the entire reaction by electron microscopy. We chose 10μM of actin to induce rapid polymerization without a lag phase. Concentrations of cofilin, coronin, and Aip1 were picked to mimic the roughly estimated ratios of these factors to actin in various cell types and tissue extracts (18,33,38,43). While many long filaments form during the first minute of assembly (Figure 1A) no polymer remains after 60 seconds in cofilin, coronin, and Aip1 (Figure 1B). Higher magnification views of the same reactions before and after treating actin filaments with the disassembly cocktail for one minute show only globular material with only a rare, very-short oligomer in a field of view (compare Figures 1C and 1D). Otherwise, the products of this disassembly reaction did not appear much different than images of the three disassembly factors alone in the absence of any actin (Figure 1E). The rapid loss of all actin filaments required all three factors because filaments were readily found after treating one minute old actin filaments with any two of three actin disassembly factors (Figure 1F). Figure 1G shows a time course of the disassembly reaction conducted in the presence of all three depolymerizers. Note that actin filaments already appear to be getting shorter within 5 seconds of adding the three disassembly factors.

**Figure 1.**
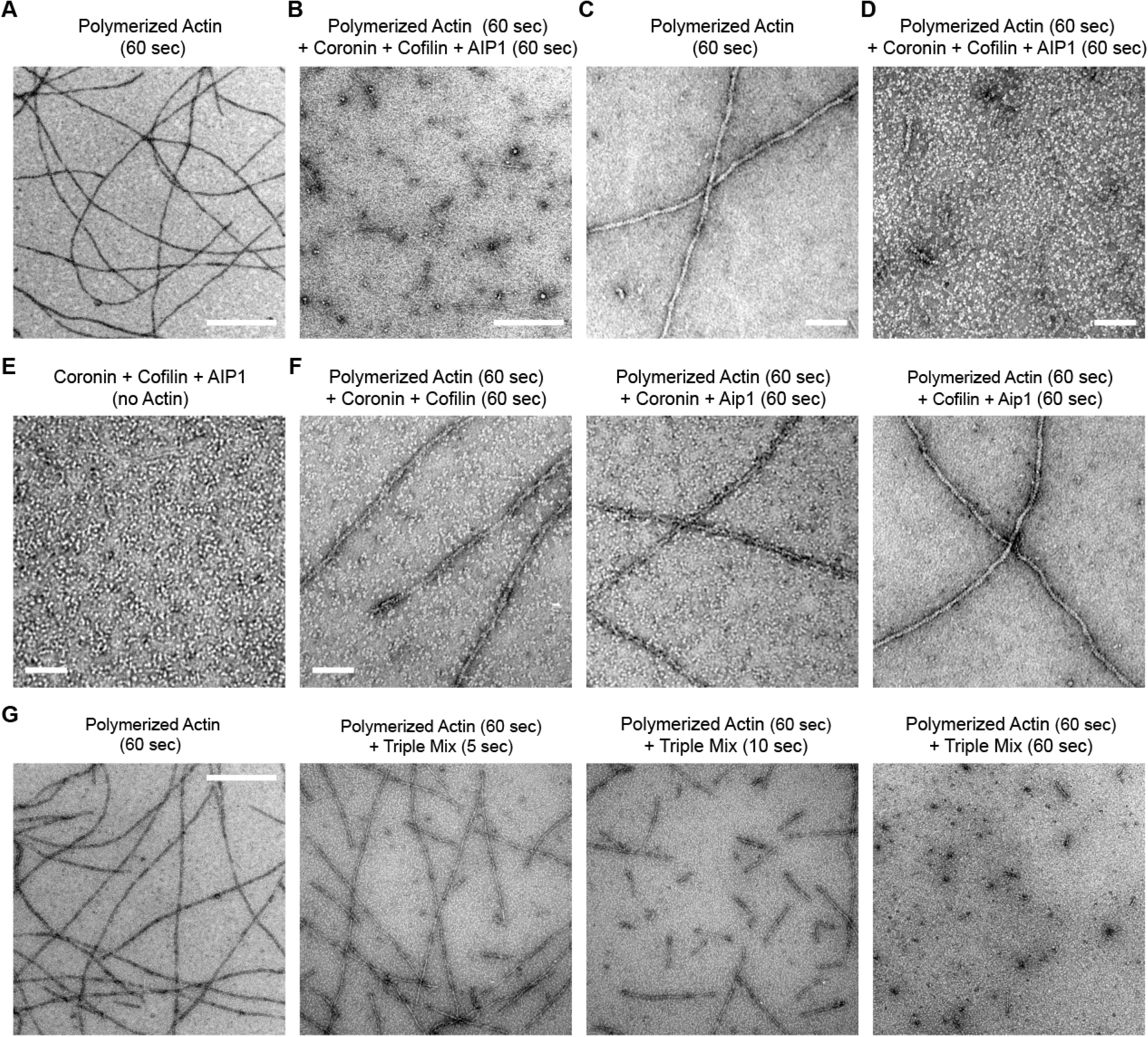
Overview of actin filament disassembly in the presence of cofilin, coronin, and Aip1 using electron microscopy. In all panels, actin is polymerized for 1 minute before any additional steps. (A) A 10 μM solution of G-actin was induced to polymerize by adding salt and ATP. The sample was fixed and prepared for visualization in the electron microscope. (B) Electron micrograph of the products produced by adding 5 cofilin, 2 coronin, and 0.2 Aip1 to filaments for 60 seconds. (C) Higher magnification image of actin filaments polymerized for 60s. (D) Higher magnification image after incubating filaments in cofilin, coronin, and Aip1 for 60s. (E) Image of coronin, cofilin, and Aip1 alone (no actin). (F) Morphology of actin filaments that were incubated for an additional 60s in the presence of various combinations of two out of the three disassembly factors. 1G) Time course of actin filament disassembly in the presence of cofilin, coronin and Aip1. The left most panel is an image of 1 minute old actin before adding the depolymerizers. Subsequent panels are images of the reaction products at various times after adding the triple mix of disassembly factors. Scale bars are 500 nm in A, B, and G and 100 nm in C-F.

We recently described a FRET-based assay to measure changes in actin polymer mass in the presence of cofilin (18). Unlike pyrene, the FRET assay is not perturbed by cofilin binding to F-actin but only reports on depolymerization. Spectroscopic methods would enable us to detect actin polymer mass that escaped detection by light or electron microscopy and would report on polymer mass at steady state in the presence of the disassembly factors.

We used the FRET assay to compare disassembly of actin as a function of three disassembly factors. A 2μM solution of actin was polymerized to steady state and mixed with various combinations of the three disassembly factors. In the presence of 1.25 μM cofilin, we observed a 30-40% decrease in polymer mass, consistent with the fact that cofilin severs actin filaments and binds ADP•actin monomers that are lost from filament ends with an affinity of 150 nM (7,44) (Figure 2A, green line). The addition of 0.1 μM Aip1 increased the initial rate of disassembly approximately five times, and the final extent of depolymerization was the same as with cofilin alone (Figure 2A, yellow line) consistent with previous results (18). When 0.75 μM coronin was added to cofilin, it led to an apparent increase in polymer mass and a decrease in the depolymerization rate which is consistent with coronin’s ability to stabilize actin filaments (41) and suppress cofilin mediated filament severing (34,36) (Figure 2A, blue line). When all three factors were combined with actin, we obtained near complete loss of polymer mass (Figure 2A, red line). Along with a rapid rate of depolymerization, the extent of disassembly in the presence of all three factors was greater than that obtained by any other combination of factors. This result indicates that the triple mix of cofilin, coronin, and Aip1 rapidly depolymerizes actin and raises the critical concentration.

**Figure 2.**
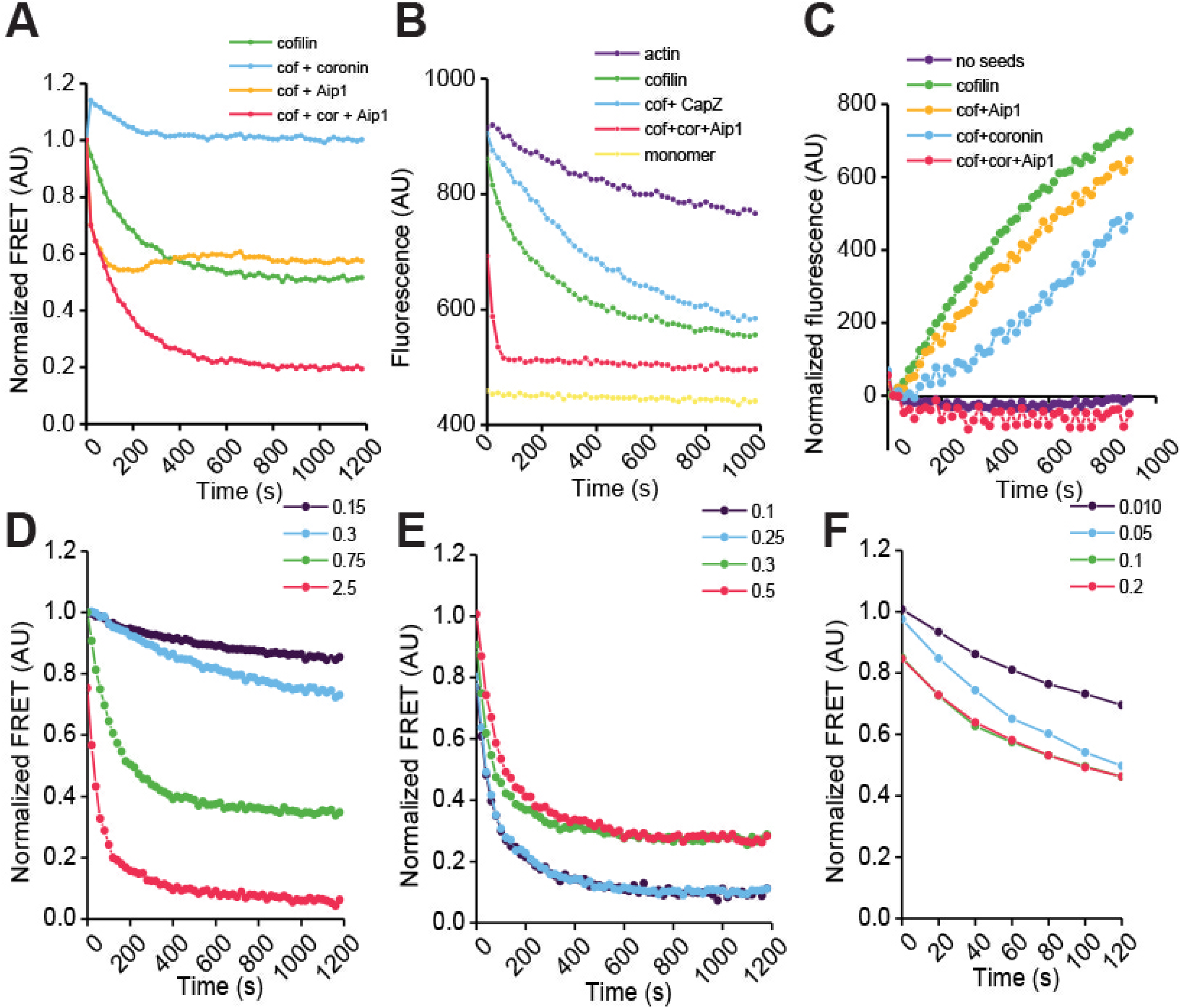
The bursting reaction proceeds with rapid kinetics to generate monomer and/or products incompetent to seed actin assembly. (A) Disassembly kinetics of 1 μM actin was monitored by loss of FRET signal in the presence of various combinations of factors. (B) Rate of monomer production was monitored by quenching of Oregon green −488 actin by Vitamin D-binding protein over time (C) Pyrene-based seeding reaction using the products of reaction (A) showed that products of the bursting reaction are not competent to seed new actin assembly. (D) Dose response curves of cofilin in the presence of 0.75 μM cofilin and 0.1 μM Aip1 shows increasing rates of disassembly with increasing amounts of cofilin (E) Lower ratios of coronin: cofilin produces more efficient disassembly whereas higher ratios may stabilize actin filaments (F) Initial rates (120 s) of disassembly with varying concentrations of Aip1 and fixed amounts of cofilin and coronin show increased rates of disassembly with increasing amounts of Aip1. Representative data from n=3 experiments is shown.

The FRET assay measures loss of actin polymer mass. We also wished to directly measure the rate of actin monomer production. We therefore developed a new monomer trap assay that reports on actin monomers. We found that Vitamin D Binding Protein (DBP), which binds exclusively to actin monomers (45,46), quenches the fluorescence of actin monomers labeled with Oregon Green (compare the purple line to the yellow line at time 0 in figure 2B). We took advantage of this property to monitor the rate of production of actin monomers in the presence of different combinations of disassembly factors. Adding DBP alone to a solution of F-actin labeled with Oregon Green leads to a slow decrease in fluorescence intensity as actin subunits are lost from the two ends of the filaments (Figure 2B purple line). Cofilin accelerated the rate of monomer production in the presence of DBP consistent with cofilin severing and the production of more ends that can shrink (Figure 2B, green line). Adding capping protein, which binds to actin filament plus ends with nanomolar affinity (47), to cofilin slowed the rate of monomer production relative to cofilin alone as expected (Figure 2B blue line). In contrast, adding coronin and Aip1 along with cofilin resulted in a very rapid production of actin monomers as indicated by the precipitous drop in Oregon Green fluorescence to its final value in less than 100 seconds (figure 2B, red line).

Although we were now convinced that the product of the disassembly reaction was largely actin monomer, to further distinguish the reaction from filament severing, we performed seeding reactions with the end products of depolymerization (Fig.2C). Severed short filaments would promote polymerization and diminish the lag phase of actin assembly. However, actin monomer or fragments with occluded ends would not serve as seeds. Unseeded pyrene actin showed a canonical lag phase prior to polymerization (Fig.2C, purple line) whereas seeds of cofilin, cofilin-coronin or cofilin-Aip1-severed F-actin filaments yielded a more effective seeding mixture consistent with the results of the FRET and electron microscopy (green, blue and yellow lines respectively). However, products of the 3x mix reaction displayed the same lag phase as monomeric actin (red line) indicating that they were incompetent to seed new actin assembly.

To gain greater insight into the mechanism by which the 3x mix of factors was able to rapidly disassemble actin with the generation of primarily actin monomer and products that were unable to seed new actin assembly, we varied the concentration of each factor in the presence of fixed concentrations of the remaining two factors. As expected, the rates of depolymerization increased with increasing concentrations of both cofilin and Aip1 (Fig.2D&F). However, coronin appeared to act best at a ratio of 5-12.5:1 cofilin: coronin and disassembly was less efficient at lower ratios (Fig.2E). This is consistent with coronin’s ability to bind and stabilize F-actin and also to bind antagonistically with cofilin (34,40,41).

The 3X mix of cofilin, coronin, and Aip1 rapidly converts actin polymer to monomer. We and others have already imaged the disassembly process using fluorescence microscopy, which also suggested a rapid dissolution of F-actin into monomers (29,34). Electron microscopy could provide additional insight into the mechanism if we could capture images of actin filaments in the process of disassembling. Working together, two people can reliably fix the sample within five seconds of adding the depolymerizers to a solution of F-actin. In figure 3A, solutions of actin were polymerized for 1 minute. Cofilin, coronin, and Aip1 were then added to these filaments and the reactions fixed after 5, 10, or 30 seconds. The low magnification images show the rapid loss of actin polymer mass over time (figure 3A, upper panels). The higher magnification images show filament architecture at these early time points (figure 3A, lower panels). Within 5 seconds of adding the triple mix, long sections of actin polymer were losing coherent structure. By ten seconds, filaments appeared shorter and were disintegrating all along their length. By 30 seconds, most of the polymer mass was gone and the few remaining filaments were short with a highly compromised, disorganized structure. A gallery of images showing dissolution of actin polymer structure in the presence of cofilin, coronin, and Aip1 is provided in supplementary figure 1.

**Figure 3.**
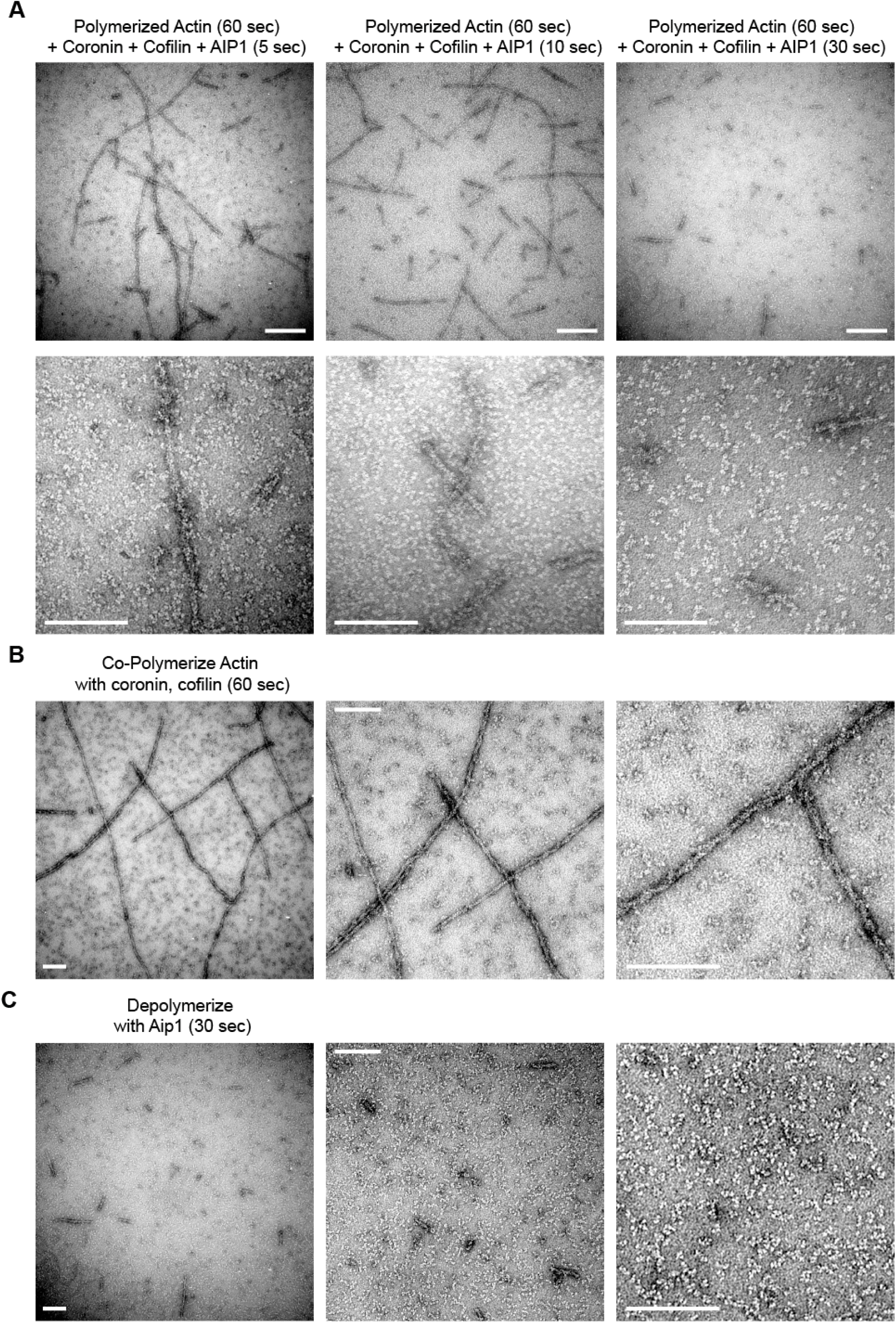
Long stretches of actin polymer disintegrate in the presence of coronin, cofilin, and Aip1. (A) Actin filaments, one minute old, were mixed with cofilin, coronin, and Aip1 and then fixed at the times indicated. The upper panels show low magnification views while the lower panels are at higher magnification. Segments of actin polymer are losing coherent structure as early as five seconds. (B) Actin filament structure is intact in the presence of cofilin and coronin alone. Each panel shows higher magnification images of actin filaments that were polymerized in the presence of cofilin and coronin for one minute. (C) Adding Aip1 to minute-old filaments copolymerized with cofilin and coronin results in the rapid disintegration of polymer into monomers and short oligomers. Each panel shows higher magnification images of the products of the reaction. Scale bars are 200 nm.

The speed of filament disintegration as well as the profound disruption of filament architecture makes it difficult to understand what is happening during the run-up to an Aip1 mediated disassembly event. Previous results with Listeria showed that the comet tails were stable in the presence of coronin and cofilin alone but disassembled upon addition of Aip1 (33). This, and the observation that coronin suppresses cofilin-mediated filament severing (34) led us to hypothesize that we could split the reaction into two steps. In the first step, we copolymerized 10 μM actin in the presence of 5 μM cofilin and 2 μM coronin and subsequently added Aip1 in a second step to disassemble these filaments. Panels in Figure 3B show three different magnifications of actin filaments that were assembled in the presence of cofilin and coronin for one minute. Panels in Figure 3C show images at three magnifications of the products of the reaction after adding Aip1 to filaments that were prepolymerized in the presence of coronin and cofilin. The filaments appeared stable in the first step (Figure 3B) and underwent dramatic disassembly within 30 seconds after treating with Aip1 (Figure 3C). Thus, at the level of filament architecture that can be assessed by electron microscopy with negative stain, the two-step reaction disassembled actin filaments through the same pathway as adding all three factors at the same time.

We examined the structure of stable actin filaments assembled in the presence of cofilin and coronin alone for 90 seconds (Figures 4A and 4B). Actin filaments can be described as a helix comprised of two protofilaments (48). In projection in the electron microscope the two protofilaments appear thin when they are on top of one another and thick when the two protofilaments are side-by-side. The distance from one thick region to the next is termed the crossover distance, which is 37 nm in pure actin and is determined by the configuration of actin subunits within the filament. The diameter of pure actin is approximately 7 nm. When actin filaments were polymerized in the presence of coronin and cofilin, we frequently found thin filaments of entirely normal twist adjacent to filaments that appeared thicker, with an altered twist and a shortened crossover distance. Figure 4B shows a representative tomogram with two filaments of distinct twists, with dots indicating the positions of crossover distances on the filaments. It is possible to distinguish individual protomers within the filament in an enlarged version of the tomogram shown in Supplementary Figure 2.

**Figure 4.**
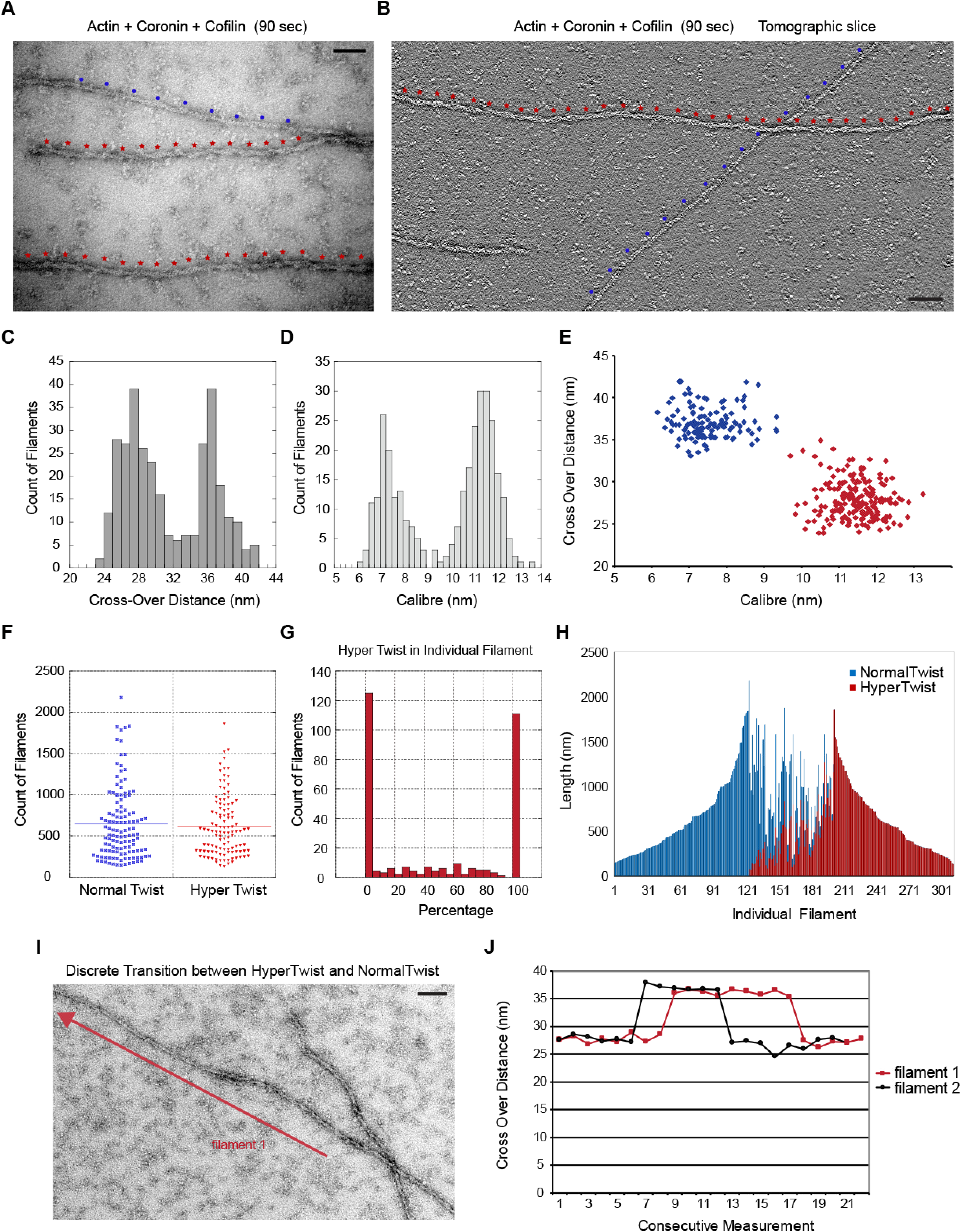
Cooperative changes in filament structure in the presence of cofilin and coronin. (A) Electron micrograph of actin filaments assembled in the presence of cofilin and coronin for 90 seconds. Red and blue dots along the filaments mark the crossover points. (B) One section of an electron tomogram showing the crossover points on two different filaments. (C) Histogram of crossover distances seen in actin filaments polymerized in the presence of cofilin and coronin for 90 seconds shows two different populations. (D) Two populations of filament widths form in the presence of cofilin and coronin. (E) Relationship between crossover distance and caliber shows that two populations of actin filaments form in the presence of cofilin and coronin.(F) Length distribution of filaments that consist either entirely of normal or hyper twist. (G) Histogram showing the percentage of hypertwist in 300 different filaments assembled in the presence of cofilin and coronin. (H) Length and portion of hypertwist of the same 300 filaments. Each bar is one filament. (I) An example of a filament containing both normal twist and hypertwist. Crossover distances were measured along the filament in the direction of the arrow and used to generate the graph for filament 1 in (J) which shows the sharp transitions between normal and hypertwisted configurations in two representative filaments. Scale bar is 25 nm in A and 50 nm in B and I.

Measuring the crossover distances over many filaments produced a bimodal distribution with one peak corresponding to ∼37 nm which is expected for pure actin and a second peak at ∼27 nm which is expected for cofilin-actin (Figure 4C). Measuring filament caliber also produced a bimodal distribution with one peak at ∼ 7 nm expected for pure actin and a second peak at ∼ 11 nm which is expected for cofilin-actin (Figure 4D). Figure 4E shows the relationship between the crossover distance and filament caliber. Filaments that are thin (∼ 7 nm) have the normal ∼ 37 nm crossover distance while filaments that are thick (∼11 – 12 nm) have a shortened crossover distance. These results are consistent with those of McGough who showed that cofilin binding to the sides of actin filaments at equilibrium alters the configuration of the filament, which can be seen with shortened crossover distance (6). Based upon figure 4E, we designated those segments of actin polymer that were thin and had a long crossover distance to be of “normal twist”. Those segments of polymer that were thick and had a shortened crossover distance were designated as “hypertwisted” due to their appearance relative to normal filaments (compare the filaments in figures 4A and 4B where the normal twist is marked with blue dots and hypertwisted segments are marked with red dots).

We used these definitions of normal twist and hypertwist to analyze the population of filaments that were formed by polymerization in the presence of cofilin and coronin. Figure 4F shows that the length distribution of filaments that are 100% hypertwisted and 100% normal twisted is the same suggesting that the two different populations of filaments have similar stabilities and that we are analyzing twist in filaments of similar length. Next, we measured crossover distances along the entire length of 300 different filaments. Figure 4G is a histogram showing the number of filaments with given percentage of hypertwist. The histogram shows a bimodal distribution where filaments are most likely 100% normal twist or 100% hypertwisted. The data in Figure 4G was replotted in Figure 4H to show the length of each filament and the relative amount of normal twist versus hypertwist. Each bar along the abscissa in figure 4H represents one filament, and the value on the ordinate is the length of that filament. Blue versus red bars indicate the portion of that filament that is normally twisted versus hypertwisted. This analysis shows that about one third of the filaments are in the normal conformation, about one-third are hypertwisted, and the remaining third of filaments of the filaments consist of co-alternating segments of normal and hyper-twist. For those filaments that contained both twists, the transition between normal and hypertwist is sharp (figure 4I and J). If cofilin was binding randomly on filaments, we would expect a single population of filaments with mixed twists. That filaments are preferentially completely saturated or entirely bare indicates that cofilin’s binding on actin is highly cooperative. Cooperative cofilin binding to aged F-actin in the ADP state has been described many times (2-8,14,16). However, calculations of expected cluster sizes based upon cofilin cooperative binding predicts only small cluster sizes (3,16). Here, we are seeing whole filaments either saturated or devoid of cofilin. In addition, the filaments in this experiment are at most only 90 seconds old, and yet half of the total polymer is in the hypertwisted state. We could not detect any hypertwisted regions in filaments polymerized in the presence of cofilin alone at these early time points (data not shown).

As cofilin preferentially binds ADP-actin filaments, we investigated coronin’s effects on phosphate release from actin as a possible explanation for coronin’s enhancement of cofilin binding. For this we used a phosphate release assay that uses the substrate 2-amino-6-mercapto-7-methylpurine ribonucleoside (42). The substrate undergoes a shift in absorbance during phosphorolysis that enables spectroscopic monitoring of the rate of phosphate release. By this assay we found that cofilin alone accelerated the rate of phosphate release from F-actin, consistent with previous results using actophorin which is the cofilin homolog in *Acanthamoeba* (25) (Figure 5A). Increasing amounts of coronin alone also accelerated the rate of phosphate release from actin (Figure 5B). Copolymerization of cofilin and coronin with actin achieved even faster rates of phosphate release than either factor on its own (Figure 5C). Some rates of phosphate release under different conditions are summarized in Figure 5D. These results show that coronin can accelerate P_i_ release from F-actin, which would help drive cofilin binding to produce the highly cooperative changes in F-actin structure on young filaments.

**Figure 5.**
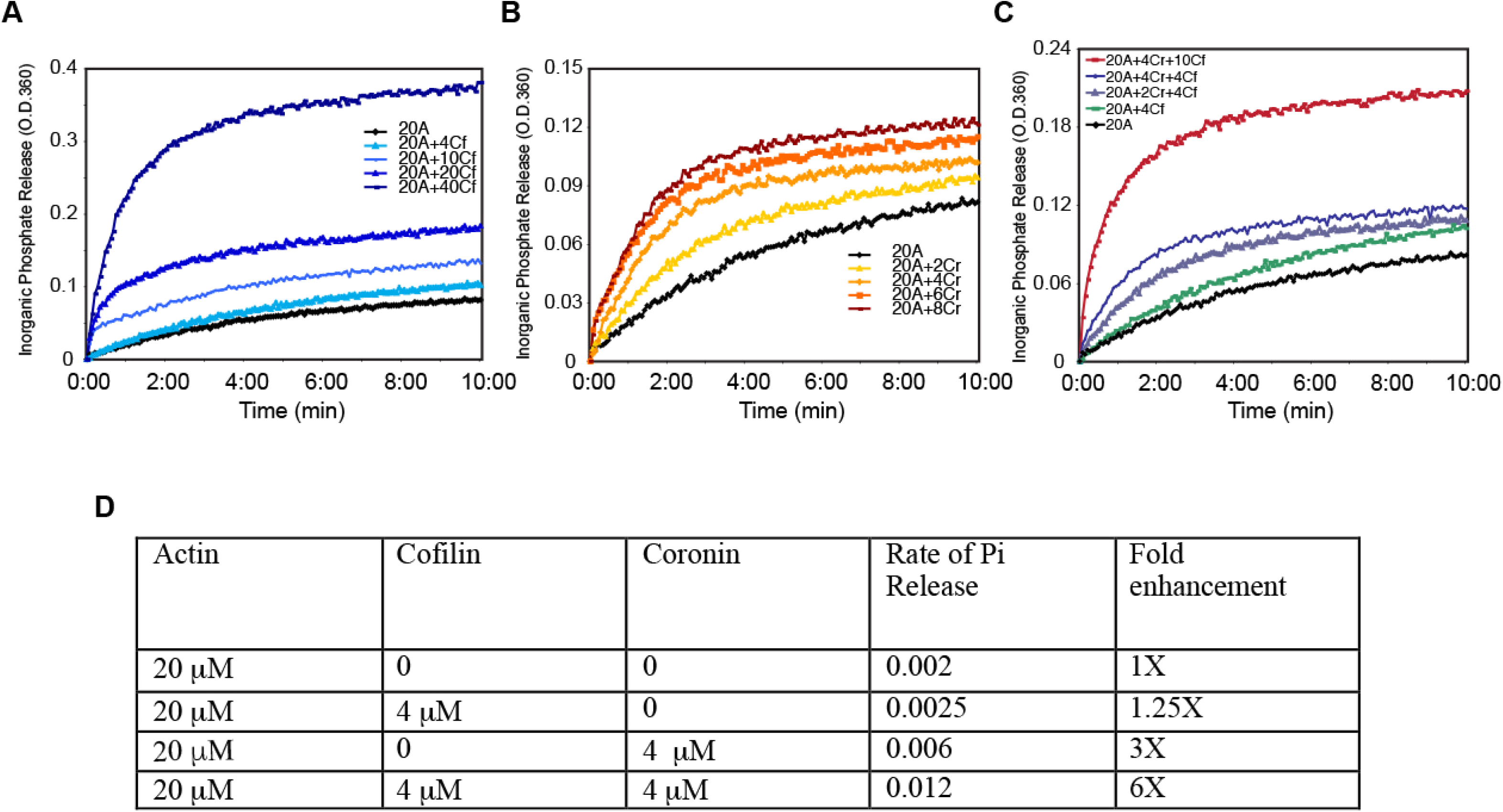
Coronin and cofilin accelerate phosphate release from F-actin. (A) Rates of phosphate release from actin filaments in the presence of increasing concentrations of cofilin alone and (B) coronin alone show accelerated phosphate release by the individual factors. (C) Together, cofilin and coronin cooperate to enhance phosphate release more than each individual factor.(D) Comparison of rates of phosphate release in the presence of actin alone, 4 μM cofilin, 4 μM coronin, or a combination of the two. Representative data from n= 4 experiments is shown.

The distribution of normal and hypertwisted sections of actin that formed as filaments were polymerizing in the presence of cofilin and coronin provided an opportunity to tests if Aip1 was disassembling actin at heterotypic junctions or along the length of hypertwisted segments. We hypothesized that it was the hypertwisted stretches of actin filaments that were visualized by EM in the presence of cofilin and coronin that underwent catastrophic disassembly in the presence of Aip1. If true, then actin filaments still visible by EM during the disassembly reaction should either have normal, 37 nm crossover distances or they should be in the process of bursting but there should be very few filaments with the shortened 27 nm crossover distance. To test this, 10 μM actin was copolymerized in the presence of 2 μM coronin and 5 μM cofilin which produced filaments with normal twist and hypertwist as expected (Fig.6A). To such a sample, 0.2 μM Aip1 was added for 5 seconds followed by rapid fixation of the filaments and visualization by EM (Fig.6B). Under these bursting conditions, in every case where we could assign a crossover distance, it was 37 nm (these filaments are marked with an N in the figure). All other sections of polymer were disintegrating as seen previously in Figure 3 with no describable order to these “polymer” segments (marked with a B in the figure). Disintegration of polymer is also evident in the gallery of images in supplemental figure 1. We were unable to detect any filaments with shortened crossovers as they underwent pronounced disassembly. These results are consistent with the hypothesis that Aip1 can target hypertwisted segments of F-actin along their length and not just heterotypic boundaries alone.

**Figure 6.**
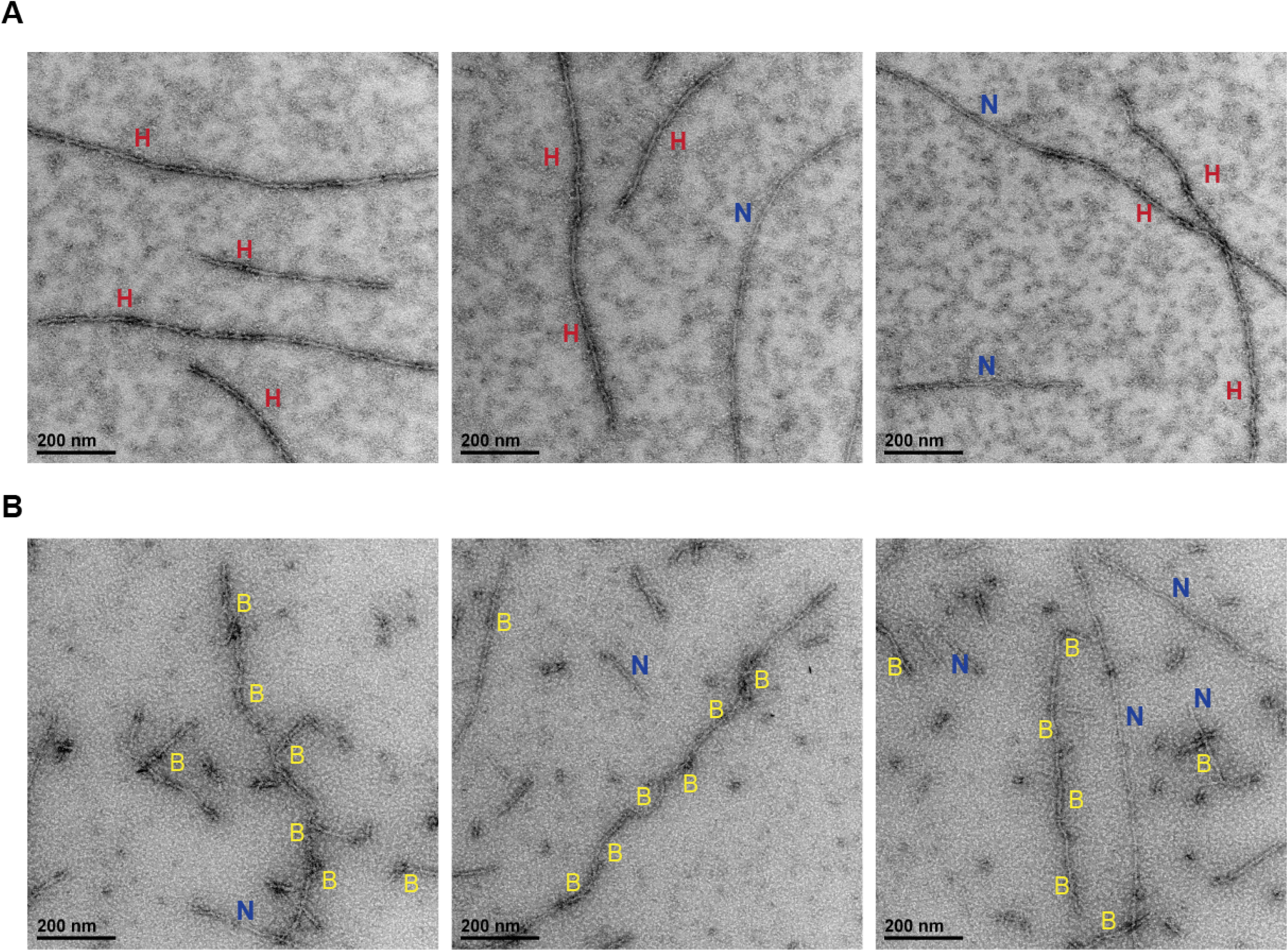
Aip1 disintegrates hypertwisted polymer. Actin was prepolymerized in the presence of coronin and cofilin in A and B for 90 seconds and then spiked with Aip1 for 5 seconds in B before fixing. 6A) Three representative images of actin filaments polymerized in the presence of coronin and cofilin for 90 seconds. Segments of polymer with normal twist are marked “N” and hypertwisted polymer is marked “H”. (B) Actin, first assembled as in 6A, five seconds after adding Aip1. “Bs” mark regions where polymer segments are disintegrating. Unmarked filaments are not obviously bursting, but crossover distances cannot be assigned. Scales bars are 200 nm.

## Discussion

We examined actin filament disassembly in the presence of cofilin, coronin, and Aip1. Our data shows that coronin accelerates the release of inorganic phosphate from F-actin to accelerate highly cooperative cofilin binding that produces long stretches of hypertwisted polymer. Aip1 disintegrates the hypertwisted sections all along their length and not just at heterotypic junctions to rapidly produce actin monomers.

The combination of coronin and cofilin produced long stretches of actin polymer with shortened crossover distance and thickened caliber indicative of cooperative cofilin binding to F-actin. In contrast, we did not see any filaments with the alternative twist or thickened caliber in the presence of cofilin alone. Multiple methods have shown that cofilin will bind cooperatively to ADP•F-actin, but not ATP or ADP•Pi F-actin. Cofilin must therefore wait for inorganic phosphate to be released from the filament, which is slow, but coronin accelerates it. Highly cooperative cofilin binding to young, newly polymerized F-actin in the presence of coronin is therefore likely to be a kinetic effect of coronin on phosphate release. Coronin locks the ATP binding cleft in a more open state (49), which would make it easier for Pi to escape the filament following ATP hydrolysis. However, the outcomes from Aip1-dependent actin disassembly are coronin-dependent. In the absence of coronin, Aip1 accelerates both severing and orderly subunit dissociation from filament ends, and the combination of Aip1 and cofilin alone has no effect the critical concentration with the barbed ends remaining open (18). In contrast, Aip1 in the presence of cofilin and coronin triggers a major disruption of filament architecture, alters the critical concentration, and caps any remaining barbed ends (33,34). Therefore, coronin contributes more to the disassembly reaction than accelerating phosphate release to promote cofilin binding.

Cofilin binds cooperatively to F-actin and alters the filament’s twist, but the exact purpose of cooperative binding for actin disassembly is still being investigated. Severing events take place at or near heterotypic junctions between cofilin-occupied and unoccupied polymer (15). Cooperative binding could contribute to cofilin severing if severing requires a cluster of contiguous cofilins (8,14). More recently, elegant work from Wioland and colleagues showed that cofilin-saturated patches of actin can depolymerize from filament barbed ends even in the presence of actin monomer and capping protein (8). Another consequence of cofilin cooperative binding is that it would allow for factors that compete with cofilin for binding to F-actin to promote severing by displacing cofilin from the filament to create a new heterotypic junction (17). Finally, earlier work showed that Aip1 disassembles actin filaments most efficiently at saturating cofilin binding densities suggesting that Aip1 might target cofilin bound polymer directly, not just the junction (18). By growing actin filaments in the presence of coronin and cofilin, we were able to obtain a population of filaments with roughly equal amounts of F-actin that were either hypertwisted or of normal twist. Electron microscopy shown here was consistent with the hypothesis that Aip1 was attacking and disintegrating hypertwisted portions of actin all along its length. Therefore, while stretches of polymer with contiguous cofilin are stable on their own, these regions appear to be targets for certain auxiliary factors to accelerate cofilin mediated filament disassembly and depolymerization.

Previous analysis of actin disassembly using live, fluorescence imaging suggested that cofilin, coronin, and Aip1 caused long segments of actin polymer to convert to monomers and very short oligomers in a single step referred to as a burst (30). The alternative view would be that bursting is simply a severing event where one of the severed fragments diffused out of the focal plane. The wholesale disruption of filament structure detected by electron microscopy and the rapid production of actin monomers detected by spectroscopy shown here are consistent with a catastrophic mode of disassembly that is distinct from filament severing. Modeling and experiment have shown that catastrophic actin filament disassembly could explain the exponential decay kinetics of Listeria actin comet tails (29). A combination of bursting and severing might also help explain the two distinct actin turnover rates detected in the actin cortex (50). In general, a catastrophic mode of filament disassembly would produce many actin monomers but few filament ends. Eliminating filaments without creating ends would explain why actin does not incorporate into the disassembling Listeria actin comet tail (29). Catastrophic elimination of filaments might be most useful when cells need to completely dissolve an actin network. In contrast, severing would produce very few monomers but many filament ends. Severing might be more useful for when cells are trying to create more actin filament barbed ends to seed rapid polymerization, as has been argued in the past (51,52). Perhaps cells can differentially regulate the activities of the disassembly factors to tune the mode of actin disassembly between severing and bursting to selectively control the morphogenesis and stability of different actin networks.

## Supplementary Figure Legends

**Supplementary Figure 1.**
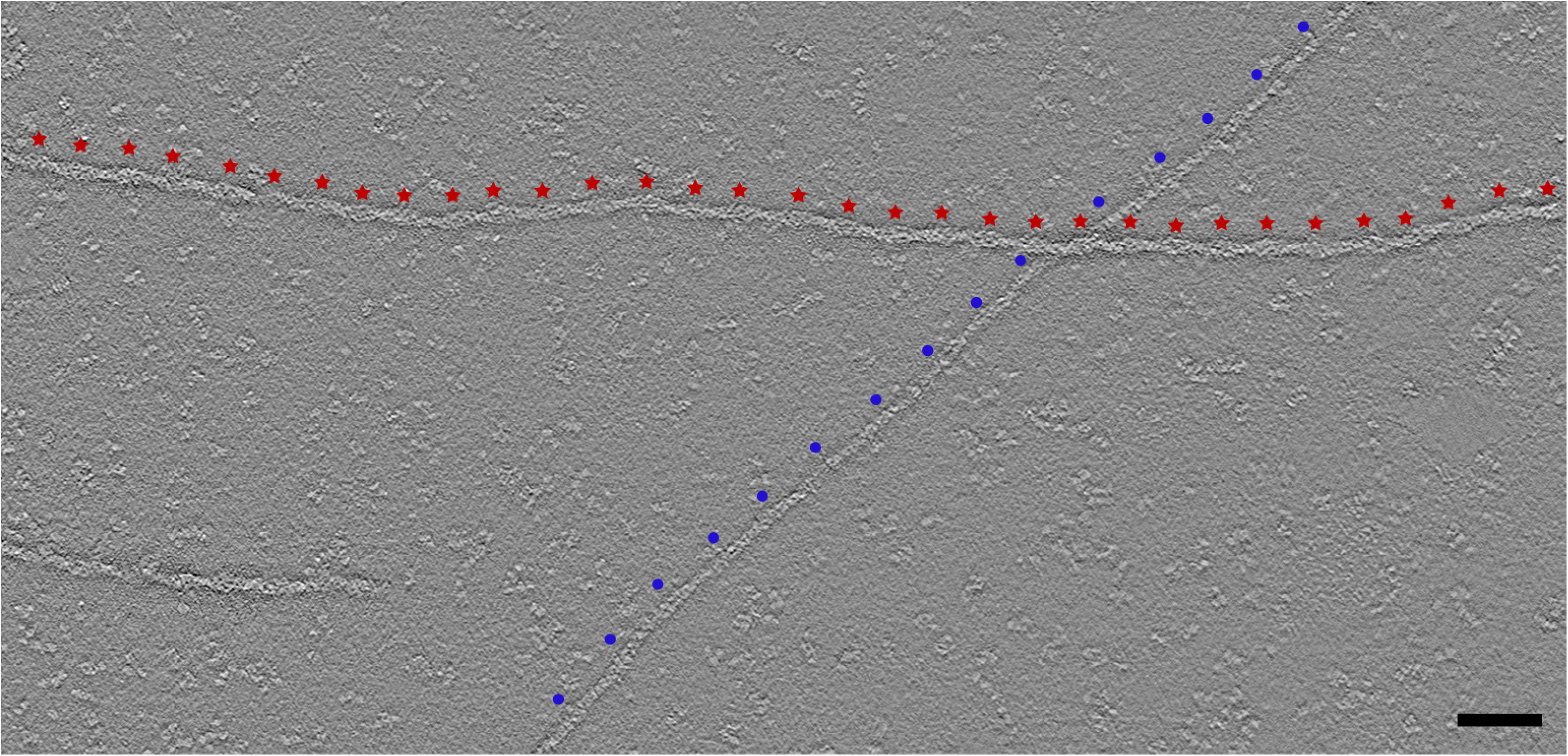
Gallery of images showing dissolution of actin filaments in the presence of cofilin, coronin, and Aip1. Actin was polymerized for 90 seconds, treated with cofilin, coronin, and Aip1 for 5 seconds, and then fixed before imaging. Filament architecture is highly distorted under these conditions as the filaments disintegrate along their length. Scale bar is 25 nm.

**Supplementary figure 2.**
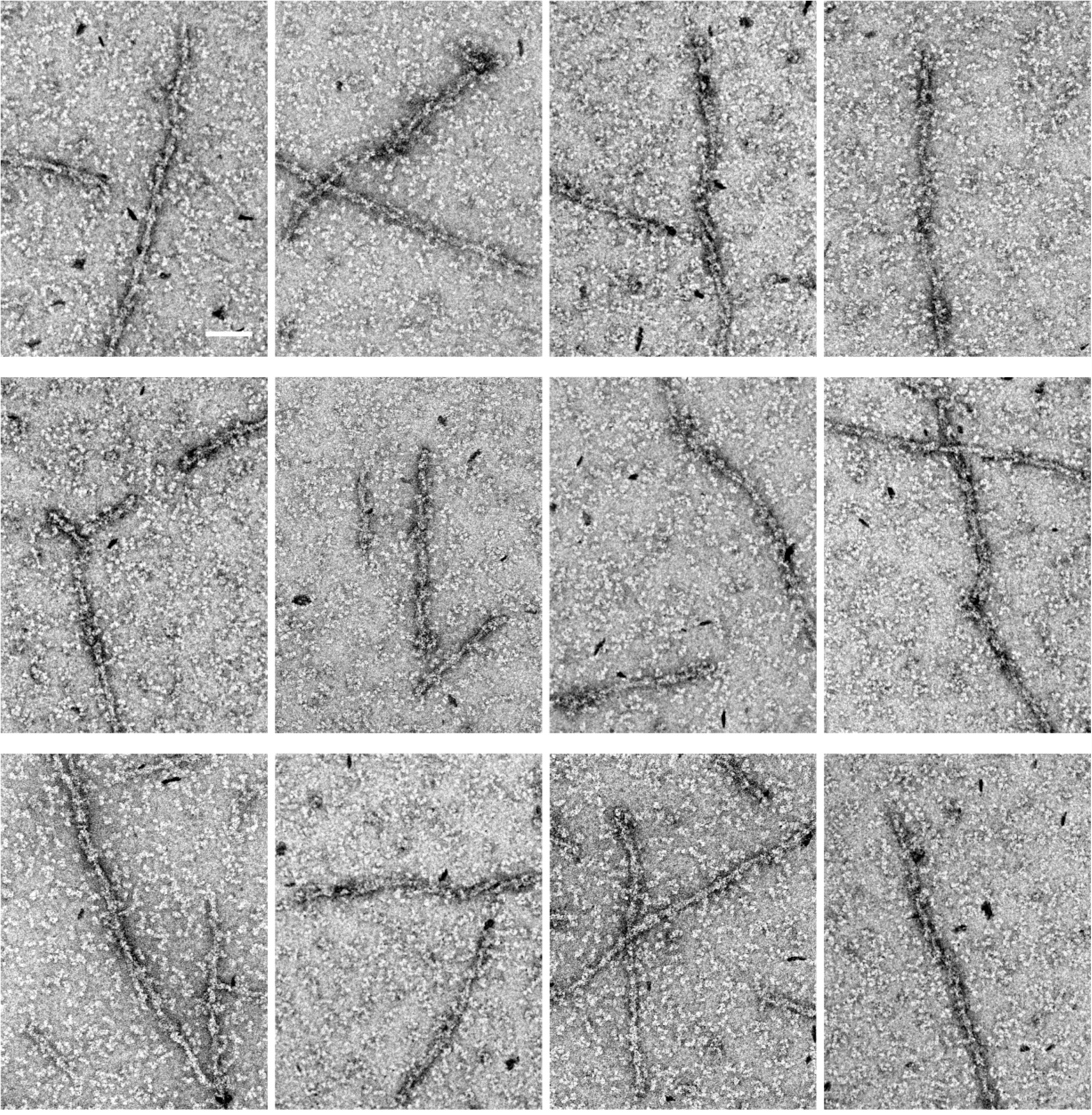
A larger image of one slice from the electron tomogram shown in figure 4B where the actin was polymerized in the presence of cofilin and coronin. Individual actin subunits are resolved in the filament lattice. Scale bar is 50 nm.

## Acknowledgements.

We thank Scott Robinson (Beckman Institute, UIUC) for use of his glow discharge unit. This work is funded by the National Institutes of Health (R01-DK098398 to V.W. Tang and R01-GM106106 to W.M. Brieher).

## Conflict of Interest Statement.

The authors declare that they have no conflicts of interest with the contents of this article.

## Author contributions.

VT and WMB performed the electron microscopy. VT performed the phosphate release assay. AVN performed the spectroscopy in figure 2 and developed the monomer trap assay. All authors analyzed results and wrote the manuscript.

## References.

Lappalainen, P., and Drubin, D. G. (1997) Cofilin promotes rapid actin filament turnover in vivo. Nature 388, 78–82

Cao, W., Goodarzi, J. P., and De La Cruz, E. M. (2006) Energetics and kinetics of cooperative cofilin-actin filament interactions. J Mol Biol 361, 257–267

De La Cruz, E. M. (2005) Cofilin binding to muscle and non-muscle actin filaments: isoform-dependent cooperative interactions. J Mol Biol 346, 557–564

Hawkins, M., Pope, B., Maciver, S. K., and Weeds, A. G. (1993) Human actin depolymerizing factor mediates a pH-sensitive destruction of actin filaments. Biochemistry 32, 9985–9993

Hayden, S. M., Miller, P. S., Brauweiler, A., and Bamburg, J. R. (1993) Analysis of the interactions of actin depolymerizing factor with G-and F-actin. Biochemistry 32, 9994– 10004

McGough, A., Pope, B., Chiu, W., and Weeds, A. (1997) Cofilin changes the twist of F-actin: implications for actin filament dynamics and cellular function. J Cell Biol 138, 771–781

Ressad, F., Didry, D., Xia, G. X., Hong, Y., Chua, N. H., Pantaloni, D., and Carlier, M. F. (1998) Kinetic analysis of the interaction of actin-depolymerizing factor (ADF)/cofilin with G- and F-actins. Comparison of plant and human ADFs and effect of phosphorylation. J Biol Chem 273, 20894–20902

Wioland, H., Guichard, B., Senju, Y., Myram, S., Lappalainen, P., Jegou, A., and Romet-Lemonne, G. (2017) ADF/Cofilin Accelerates Actin Dynamics by Severing Filaments and Promoting Their Depolymerization at Both Ends. Curr Biol

Galkin, V. E., Orlova, A., Kudryashov, D. S., Solodukhin, A., Reisler, E., Schroder, G. F., and Egelman, E. H. (2011) Remodeling of actin filaments by ADF/cofilin proteins. Proc Natl Acad Sci U S A 108, 20568–20572

McCullough, B. R., Blanchoin, L., Martiel, J. L., and De la Cruz, E. M. (2008) Cofilin increases the bending flexibility of actin filaments: implications for severing and cell mechanics. J Mol Biol 381, 550–558

Prochniewicz, E., Janson, N., Thomas, D. D., and De la Cruz, E. M. (2005) Cofilin increases the torsional flexibility and dynamics of actin filaments. J Mol Biol 353, 990– 1000

De La Cruz, E. M., Martiel, J. L., and Blanchoin, L. (2015) Mechanical heterogeneity favors fragmentation of strained actin filaments. Biophys J 108, 2270–2281

Michelot, A., Grassart, A., Okreglak, V., Costanzo, M., Boone, C., and Drubin, D. G. (2013) Actin filament elongation in Arp2/3-derived networks is controlled by three distinct mechanisms. Dev Cell 24, 182–195

Ngo, K. X., Kodera, N., Katayama, E., Ando, T., and Uyeda, T. Q. (2015) Cofilin-induced unidirectional cooperative conformational changes in actin filaments revealed by high-speed atomic force microscopy. Elife 4

Suarez, C., Roland, J., Boujemaa-Paterski, R., Kang, H., McCullough, B. R., Reymann, A. C., Guerin, C., Martiel, J. L., De la Cruz, E. M., and Blanchoin, L. (2011) Cofilin tunes the nucleotide state of actin filaments and severs at bare and decorated segment boundaries. Curr Biol 21, 862–868

Andrianantoandro, E., and Pollard, T. D. (2006) Mechanism of actin filament turnover by severing and nucleation at different concentrations of ADF/cofilin. Mol Cell 24, 13–23

Elam, W. A., Kang, H., and De La Cruz, E. M. (2013) Competitive displacement of cofilin can promote actin filament severing. Biochem Biophys Res Commun 438, 728–731

Nadkarni, A. V., and Brieher, W. M. (2014) Aip1 destabilizes cofilin-saturated actin filaments by severing and accelerating monomer dissociation from ends. Curr Biol 24, 2749–2757

Moriyama, K., and Yahara, I. (1999) Two activities of cofilin, severing and accelerating directional depolymerization of actin filaments, are affected differentially by mutations around the actin-binding helix. EMBO J 18, 6752–6761

Brieher, W. (2013) Mechanisms of actin disassembly. Mol Biol Cell 24, 2299–2302

Clark, M. G., and Amberg, D. C. (2007) Biochemical and genetic analyses provide insight into the structural and mechanistic properties of actin filament disassembly by the Aip1p cofilin complex in Saccharomyces cerevisiae. Genetics 176, 1527–1539

Clark, M. G., Teply, J., Haarer, B. K., Viggiano, S. C., Sept, D., and Amberg, D. C. (2006) A genetic dissection of Aip1p’s interactions leads to a model for Aip1p-cofilin cooperative activities. Mol Biol Cell 17, 1971–1984

Nomura, K., Hayakawa, K., Tatsumi, H., and Ono, S. (2016) Actin-interacting Protein 1 Promotes Disassembly of Actin-depolymerizing Factor/Cofilin-bound Actin Filaments in a pH-dependent Manner. J Biol Chem 291, 5146–5156

Blanchoin, L., and Pollard, T. D. (2002) Hydrolysis of ATP by polymerized actin depends on the bound divalent cation but not profilin. Biochemistry 41, 597–602

Blanchoin, L., and Pollard, T. D. (1999) Mechanism of interaction of Acanthamoeba actophorin (ADF/Cofilin) with actin filaments. J Biol Chem 274, 15538–15546

Carlier, M. F. (1987) Measurement of Pi dissociation from actin filaments following ATP hydrolysis using a linked enzyme assay. Biochem Biophys Res Commun 143, 1069–1075

Carlier, M. F., and Pantaloni, D. (1986) Direct evidence for ADP-Pi-F-actin as the major intermediate in ATP-actin polymerization. Rate of dissociation of Pi from actin filaments. Biochemistry 25, 7789–7792

Carlier, M. F., Laurent, V., Santolini, J., Melki, R., Didry, D., Xia, G. X., Hong, Y., Chua, N. H., and Pantaloni, D. (1997) Actin depolymerizing factor (ADF/cofilin) enhances the rate of filament turnover: implication in actin-based motility. J Cell Biol 136, 1307–1322

Kueh, H. Y., Brieher, W. M., and Mitchison, T. J. (2010) Quantitative analysis of actin turnover in Listeria comet tails: evidence for catastrophic filament turnover. Biophys J 99, 2153–2162

Kueh, H. Y., Charras, G. T., Mitchison, T. J., and Brieher, W. M. (2008) Actin disassembly by cofilin, coronin, and Aip1 occurs in bursts and is inhibited by barbed-end cappers. J Cell Biol 182, 341–353

Theriot, J. A., and Mitchison, T. J. (1991) Actin microfilament dynamics in locomoting cells. Nature 352, 126–131

Theriot, J. A., Mitchison, T. J., Tilney, L. G., and Portnoy, D. A. (1992) The rate of actin-based motility of intracellular Listeria monocytogenes equals the rate of actin polymerization. Nature 357, 257–260

Brieher, W. M., Kueh, H. Y., Ballif, B. A., and Mitchison, T. J. (2006) Rapid actin monomer-insensitive depolymerization of Listeria actin comet tails by cofilin, coronin, and Aip1. J Cell Biol 175, 315–324

Jansen, S., Collins, A., Chin, S. M., Ydenberg, C. A., Gelles, J., and Goode, B. L. (2015) Single-molecule imaging of a three-component ordered actin disassembly mechanism. Nat Commun 6, 7202

de Hostos, E. L., Bradtke, B., Lottspeich, F., Guggenheim, R., and Gerisch, G. (1991) Coronin, an actin binding protein of Dictyostelium discoideum localized to cell surface projections, has sequence similarities to G protein beta subunits. EMBO J 10, 4097–4104

Gandhi, M., Achard, V., Blanchoin, L., and Goode, B. L. (2009) Coronin switches roles in actin disassembly depending on the nucleotide state of actin. Mol Cell 34, 364–374

Mikati, M. A., Breitsprecher, D., Jansen, S., Reisler, E., and Goode, B. L. (2015) Coronin Enhances Actin Filament Severing by Recruiting Cofilin to Filament Sides and Altering F-Actin Conformation. J Mol Biol 427, 3137–3147

Rodal, A. A., Tetreault, J. W., Lappalainen, P., Drubin, D. G., and Amberg, D. C. (1999) Aip1p interacts with cofilin to disassemble actin filaments. J Cell Biol 145, 1251–1264

Okreglak, V., and Drubin, D. G. (2010) Loss of Aip1 reveals a role in maintaining the actin monomer pool and an in vivo oligomer assembly pathway. J Cell Biol 188, 769–777

Cai, L., Makhov, A. M., and Bear, J. E. (2007) F-actin binding is essential for coronin 1B function in vivo. J Cell Sci 120, 1779–1790

Galkin, V. E., Orlova, A., Brieher, W., Kueh, H. Y., Mitchison, T. J., and Egelman, E. H. (2008) Coronin-1A stabilizes F-actin by bridging adjacent actin protomers and stapling opposite strands of the actin filament. J Mol Biol 376, 607–613

Webb, M. R. (1992) A continuous spectrophotometric assay for inorganic phosphate and for measuring phosphate release kinetics in biological systems. Proc Natl Acad Sci U S A 89, 4884–4887

Goode, B. L., Wong, J. J., Butty, A. C., Peter, M., McCormack, A. L., Yates, J. R., Drubin, D. G., and Barnes, G. (1999) Coronin promotes the rapid assembly and cross-linking of actin filaments and may link the actin and microtubule cytoskeletons in yeast. J Cell Biol 144, 83–98

Blanchoin, L., and Pollard, T. D. (1998) Interaction of actin monomers with Acanthamoeba actophorin (ADF/cofilin) and profilin. J Biol Chem 273, 25106–25111

Lees, A., Haddad, J. G., and Lin, S. (1984) Brevin and vitamin D binding protein: comparison of the effects of two serum proteins on actin assembly and disassembly. Biochemistry 23, 3038–3047

Van Baelen, H., Bouillon, R., and De Moor, P. (1980) Vitamin D-binding protein (Gc-globulin) binds actin. J Biol Chem 255, 2270–2272

Wear, M. A., Yamashita, A., Kim, K., Maeda, Y., and Cooper, J. A. (2003) How capping protein binds the barbed end of the actin filament. Curr Biol 13, 1531–1537

Howard, J. (2001) Mechanics of Motor Proteins and the Cytoskeleton, Sinauer Associates

Ge, P., Durer, Z. A., Kudryashov, D., Zhou, Z. H., and Reisler, E. (2014) Cryo-EM reveals different coronin binding modes for ADP- and ADP-BeFx actin filaments. Nat Struct Mol Biol 21, 1075–1081

Fritzsche, M., Lewalle, A., Duke, T., Kruse, K., and Charras, G. (2013) Analysis of turnover dynamics of the submembranous actin cortex. Mol Biol Cell 24, 757–767

Bravo-Cordero, J. J., Magalhaes, M. A., Eddy, R. J., Hodgson, L., and Condeelis, J. (2013) Functions of cofilin in cell locomotion and invasion. Nat Rev Mol Cell Biol 14, 405–415

Ghosh, M., Song, X., Mouneimne, G., Sidani, M., Lawrence, D. S., and Condeelis, J. S. (2004) Cofilin promotes actin polymerization and defines the direction of cell motility. Science 304, 743–746

